# A multi-scale structural and biophysical atlas of TCR-peptide-HLA recognition dynamics

**DOI:** 10.64898/2026.08.19.745737

**Authors:** Shan Zhang, Yuzhou Long, Taizhong Wang, Qinglu Zhong, Jinhui Li, Lei Fu

## Abstract

Dynamic interactions between T cell receptor (TCR) and peptide-human leukocyte antigen (pHLA) complexes are central to peptide-specific immune recognition, influencing T cell activation and immune responses. While structural biology has provided valuable static structures of TCR-pHLA complexes, systematic datasets capturing their dynamic and interaction patterns remain limited. Here, we present DynaTPH, a curated structural dynamics dataset of human TCR-pHLA complexes. DynaTPH integrates TCR-pHLA structures, covering both HLA class I and class II complexes, and extends these static structural resources with standardized molecular dynamics simulations and derived biophysical properties. Through a multi-stage filtering procedure, we identified 256 representative complexes and performed standardized all-atom molecular dynamics simulations for each system, corresponding to a cumulative simulation time of 38.4 μs. The dataset includes static structures, trajectories, corresponding frames, and derived physicochemical properties, including hydrogen bonds, intermolecular contacts, solvent accessibility, and backbone flexibility. By capturing the conformational flexibility and dynamic interaction patterns across diverse TCR-pHLA interfaces, DynaTPH extends static structural resources with multidimensional biophysical information. This dataset enables systematic investigation of TCR-pHLA recognition dynamics and supports applications in TCR engineering, vaccine design, and immune tolerance research and artificial intelligence-driven computational immunology.

## Background & Summary

The co-recognition of peptides presented by major histocompatibility complex (MHC) molecules by T cell receptors (TCRs) underlies antigen-specific adaptive immune recognition. In humans, this process is mediated through TCR engagement with peptides bound to human leukocyte antigen (HLA) molecules^1–3^. Peptides are presented on the cell surface by HLA molecules and subsequently recognized by TCRs, forming TCR-pHLA interactions^2^. These molecular interaction enables the immune system to distinguish infected or abnormal cells from healthy ones, thereby initiating appropriate immune responses^3,4^. The specificity and affinity of TCR-pHLA binding govern T cell activation and shape immune responses against pathogens, tumors, and self-antigens. This recognition disorder is associated with various diseases, including cancer^5^, autoimmune diseases^6^, and transplant rejection^1^. Consequently, TCR-pHLA interactions are crucial for the development of immunotherapies, vaccines, and antigen-specific tolerance strategies^7–9^.

Peptide-presenting HLA molecules are classified into class I and class II, a distinction that is likewise reflected in TCR-pHLA complexes and is accompanied by pronounced structural and functional differences. Class I molecules include the classical HLA-A, HLA-B, and HLA-C, as well as non-classical members such as HLA-E. The peptide-binding groove in HLA class I is formed by the α1 and α2 domains of the heavy chain^10,11^. This groove is relatively narrow and closed at both termini, allowing it to bind shorter peptides. These peptides are typically derived from endogenous proteins and are recognized by CD8^+^ T cells^11,12^. In contrast, class II molecules, such as HLA-DR, HLA-DP, and HLA-DQ, form their peptide-binding groove through the combination of the α and β chains^11^. The groove is more open and wider, enabling the binding of longer and more flexible peptides. These peptides are generally exogenous and are recognized by CD4⁺ T cells^11–13^.

Despite the substantial biomedical importance of TCR-pHLA recognition, its molecular mechanisms remain incompletely understood. Public databases such as IEDB^14^, VDJdb^15^, and TCR3D^16^ provide extensive sequence-level information, and some entries are linked to experimentally resolved crystal or cryo-electron microscopy structures. These structural resources have greatly facilitated the systematic organization and analysis of TCR-pHLA complexes and have provided important insights into their overall binding geometries. However, such experimentally determined structures primarily represent static conformational states and are therefore limited in their ability to capture the dynamic features underlying molecular recognition. TCR-pHLA interactions are highly complex and dynamic processes that begin with peptide presentation by HLA molecules and proceed through TCR recognition of the resulting pHLA complexes^2^. These interactions involve conformational fluctuations, local flexibility, and transient intermolecular contacts, all of which contribute to antigen specificity, cross-reactivity, and downstream signaling^17^. Consequently, important dynamic properties of TCR-pHLA complexes cannot be fully characterized from single static structures alone. These limitations highlight the need for standardized datasets that systematically describe the dynamic behavior of TCR-pHLA complexes.

Molecular dynamics (MD) simulations provide an effective strategy for extending static structural information into dynamic conformational landscapes. By capturing the time-resolved behavior of biomolecular systems at atomic resolution, MD enables the characterization of conformational flexibility, intermolecular interactions, and transient binding states under physiologically relevant conditions^18,19^. Such dynamic information is essential not only for advancing mechanistic understanding of immune recognition, but also for providing structural data resources that may support the development of computational models for TCR-pHLA interactions. At present, most artificial intelligence methods for predicting TCR-pHLA interactions^20,21^ or designing TCR mimics^22–24^ rely primarily on sequence features and static structural representations, which may not fully capture conformational rearrangements and dynamic contact formation during molecular recognition. Accordingly, MD-derived datasets are increasingly recognized as an important resource for bridging static structural information with functional dynamics. MD simulations not only provide trajectories that capture conformational changes of proteins, but also enable the extraction of quantitative biophysical descriptors that characterize molecular recognition at multiple levels. These descriptors, such as hydrogen bond, motions of protein residues, solvent accessibility, and conformational flexibility, among other biophysical features, provide interpretable representations of protein dynamics and binding mechanisms that cannot be directly obtained from static structures. Such dynamic structural information and derived biophysical features can complement sequence and static structure based representations in artificial intelligence models, offering opportunities to develop more physics-informed approaches for molecular recognition prediction and biomolecular design. Recent studies have shown that incorporating protein dynamic data and physicochemical properties into artificial intelligence frameworks can improve the interpretability and predictive performance of models in protein related tasks^25,26^. Several MD datasets for biomolecular systems already exist, such as the MemProD^27^ platform and GPCRmd^28^ for membrane proteins, and mdCATH^29^ for protein dynamics. However, to date, a comprehensive and systematically curated MD dataset specifically focused on TCR-pHLA complexes remains unavailable.

To address this gap, we present DynaTPH, a high-quality MD simulation dataset that covers a broad range of publicly available human TCR-pHLA complex structures, including complexes with both HLA class I and class II molecules, thereby filling a gap in dynamic structural data for these systems. We first integrated records from six major public TCR-HLA repositories, comprising more than 700,000 entries in total. After systematic screening and filtering, we identified 256 non-redundant human TCR-pHLA complexes with complete and reliable structural information. We then curated these structures to remove incomplete complexes or those with non-canonical binding geometries and to resolve inconsistencies in chain annotation and oligomeric state. Each curated complex was subjected to standardized MD simulations under uniform protocols to capture conformational dynamics and interfacial interaction patterns. Equilibrated trajectory segments were subsequently extracted for downstream analyses to derive multidimensional biophysical descriptors.

The DynaTPH dataset provides a comprehensive resource for investigating TCR-pHLA interactions within a dynamic structural framework, integrating static structures, molecular dynamics trajectories, and derived physicochemical properties organized across multiple levels, including peptide level, ternary complex level, and interface level. These data capture complementary aspects of TCR-pHLA recognition, from static conformations and time-resolved structural dynamics to quantitative molecular interaction features. By integrating these multiple layers of information within a unified framework, DynaTPH enables systematic characterization of TCR-pHLA interactions across diverse complexes. This integrated representation links sequence, structure, and dynamic interaction information and may facilitate data-driven analyses such as antigen prioritization, TCR engineering, and mechanistic studies of immune recognition. More broadly, DynaTPH provides a foundational resource for future studies of antigen recognition and may support the development of neoantigen-targeted immunotherapies and personalized vaccine design.

## Methods

The construction of the DynaTPH database comprises three key stages: data collection and screening, structure filtering and preprocessing, and MD simulations and multidimensional physicochemical data generation. The overall flowchart is illustrated in Fig. 1.

**Fig. 1.**
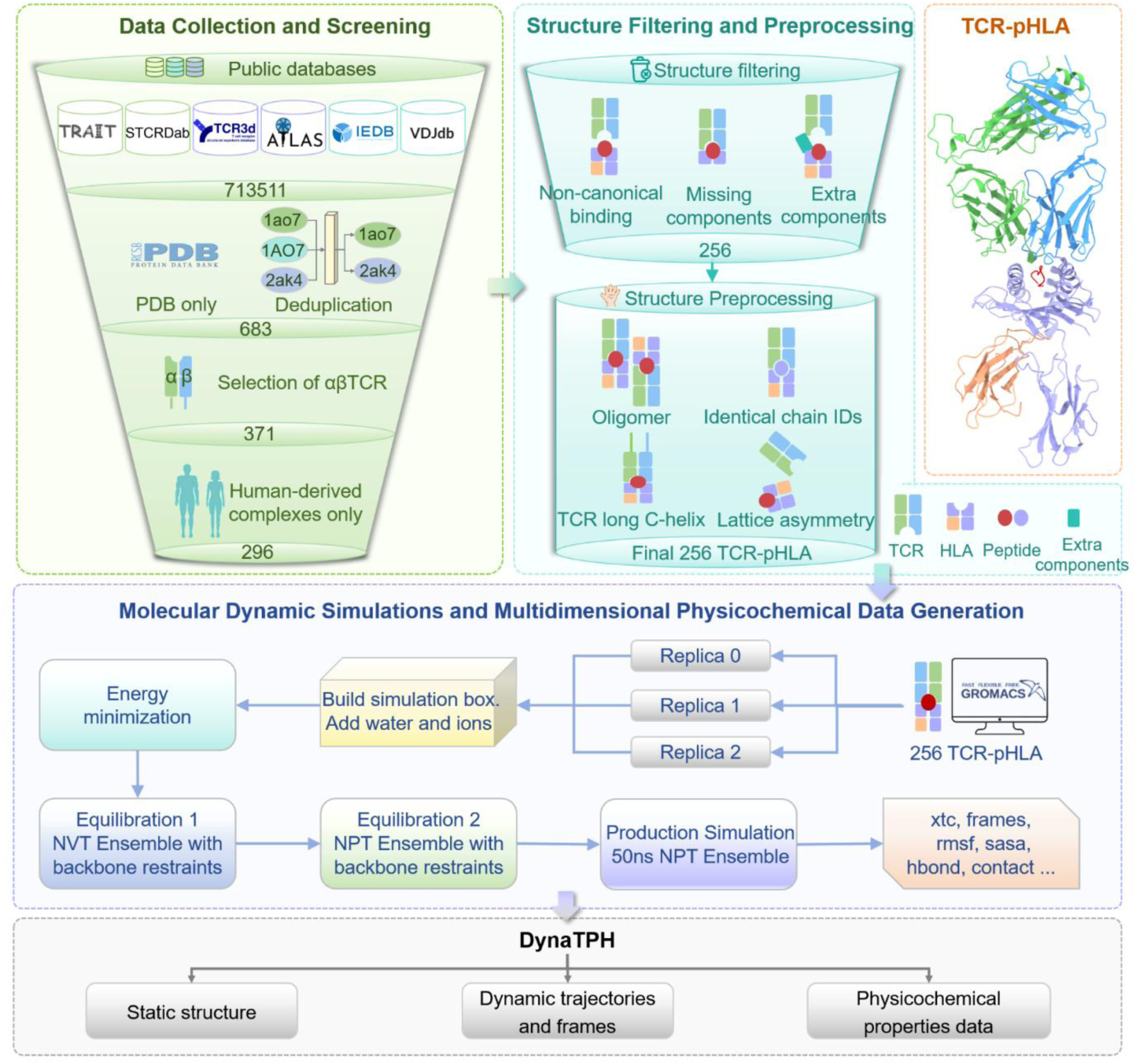
Flowchart for the construction of the DynaTPH database. The example TCR-pHLA structure is PDB: 1BD2 (HLA class I).

### Data collection and screening

To define the data sources for dataset construction, we systematically reviewed 10 publicly available databases and resources related to TCR-pHLA interactions (Table S1). The reviewed resources were compared based on their coverage of TCR sequences, antigen specificity annotations, HLA information, and structural information. Six databases were subsequently selected for data integration because they either directly provide experimentally determined TCR-pHLA structures or contain sufficient sequence and annotation information to facilitate the identification and reconstruction of corresponding structural complexes. These resources included IEDB^14^, VDJdb^15^, TCR3d^16^, TRAIT^30^, STCRDab^31^ and ATLAS^32^, yielding a total of 713,511 TCR-pMHC-related entries (Fig. 2).

**Fig. 2.**
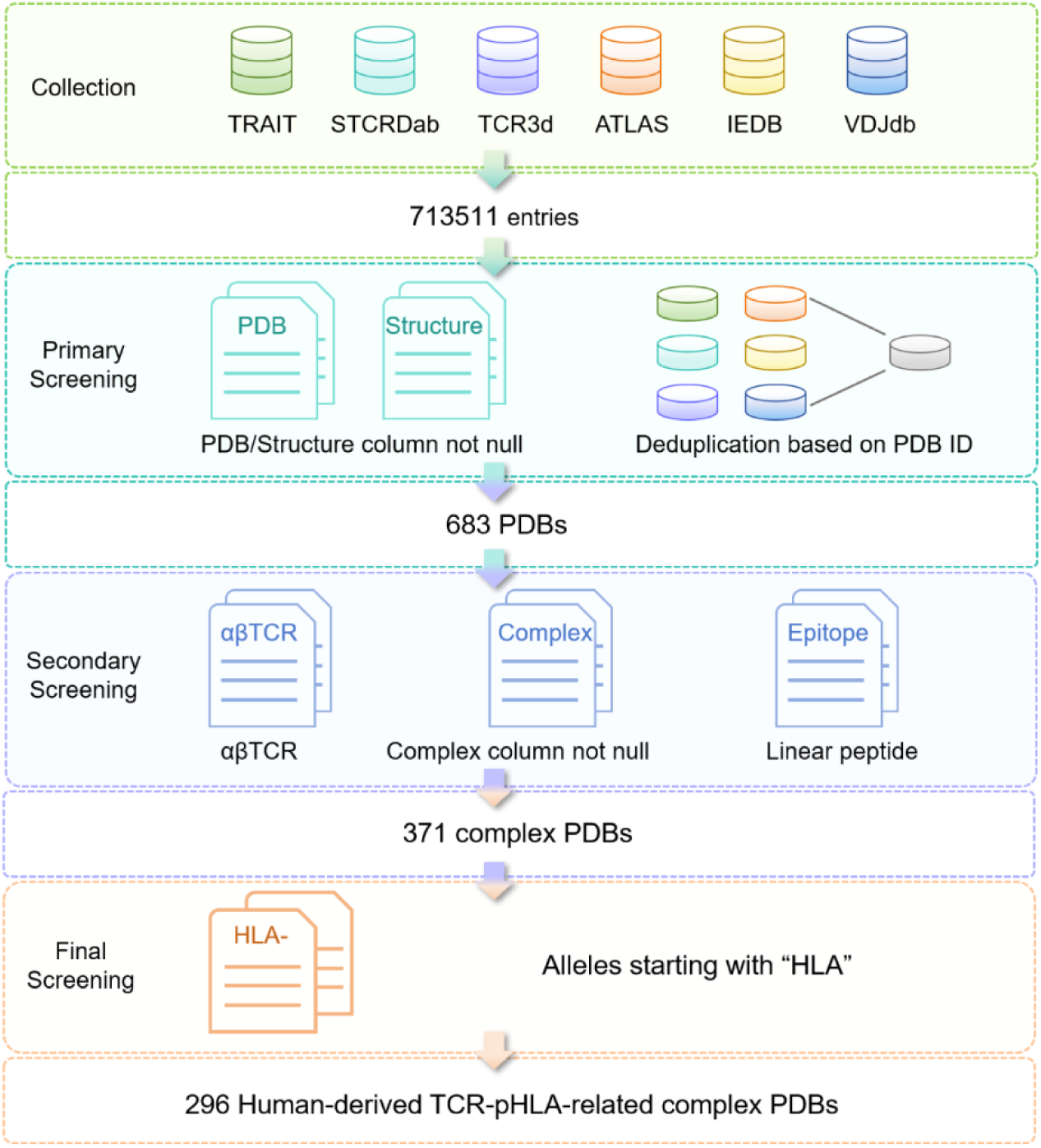
Schematic diagram of the data collection and screening process. The standards and results applied for each screening step were displayed, resulting in 296 human-derived structures for structure filtering and correction in the next step.

In the primary screening, entries with available PDB structures were selected from each database and duplicates were removed. Based on these criteria, 683 entries were retained.

Subsequently, in the secondary screening, entries were further filtered based on database-specific annotations, including complex type, peptide characteristics, and TCR chain information where available. Only entries corresponding to TCR-pMHC complexes were retained. Complexes containing linear peptide antigens were selected to ensure biological relevance. For databases providing TCR chain annotations, αβTCR complexes were preferentially retained, as αβTCRs represent the primary receptors responsible for recognizing peptide antigens presented by HLA molecules^3,12,33,34^. Because annotation formats and completeness varied across databases, these criteria were applied in a database-dependent manner. This step yielded 371 PDB entries for further processing.

Finally, only human-derived complexes were retained. Structures associated with HLA molecules were selected based on database annotations and structure nomenclature, excluding mouse and other non-human MHC complexes. This yielded 296 unique PDB IDs containing both structural and sequence information for human TCR-pHLA-related complexes.

### Structure filtering and preprocessing

A total of 296 corresponding structures were retrieved from the RCSB Protein Data Bank (https://www.rcsb.org/)^35^. Each PDB structure was systematically inspected to assess structural completeness, assembly quality, and binding interface integrity. Consisting of two steps, filter out structures that do not meet the TCR-pHLA geometric standards (Structure filtering), and then perform structural preprocessing on the retained complexes (Structure preprocessing) to generate standardized initial structures for MD simulation.

### Structure filtering

All 296 structures underwent structural quality control to assess whether they represented properly assembled TCR-pHLA complexes. The filtering criteria required the presence of all three components of the ternary complex (TCR, peptide, and HLA), two well-defined binding interfaces (peptide-HLA, TCR-pHLA), and confirmation of αβTCR complex identity. Structures that did not meet these criteria were excluded. The major excluded categories comprised three types of complexes, with representative examples shown in Fig. 1 and Fig. S1. First, non-canonical binding assemblies were removed when the TCR and pHLA were spatially separated and unlikely to maintain stable interactions. Second, incomplete complexes that lacking the peptide, the HLA component, or essential residues at the binding interface were excluded. This step excluded entries containing only two components or disrupted ternary interfaces. Third, complexes containing extra molecular components beyond the TCR-pHLA ternary assembly were excluded. After structural quality control, 40 entries were removed, resulting in a final dataset of 256 PDB structures (Fig. 1 and Table 1).

**Table 1.**
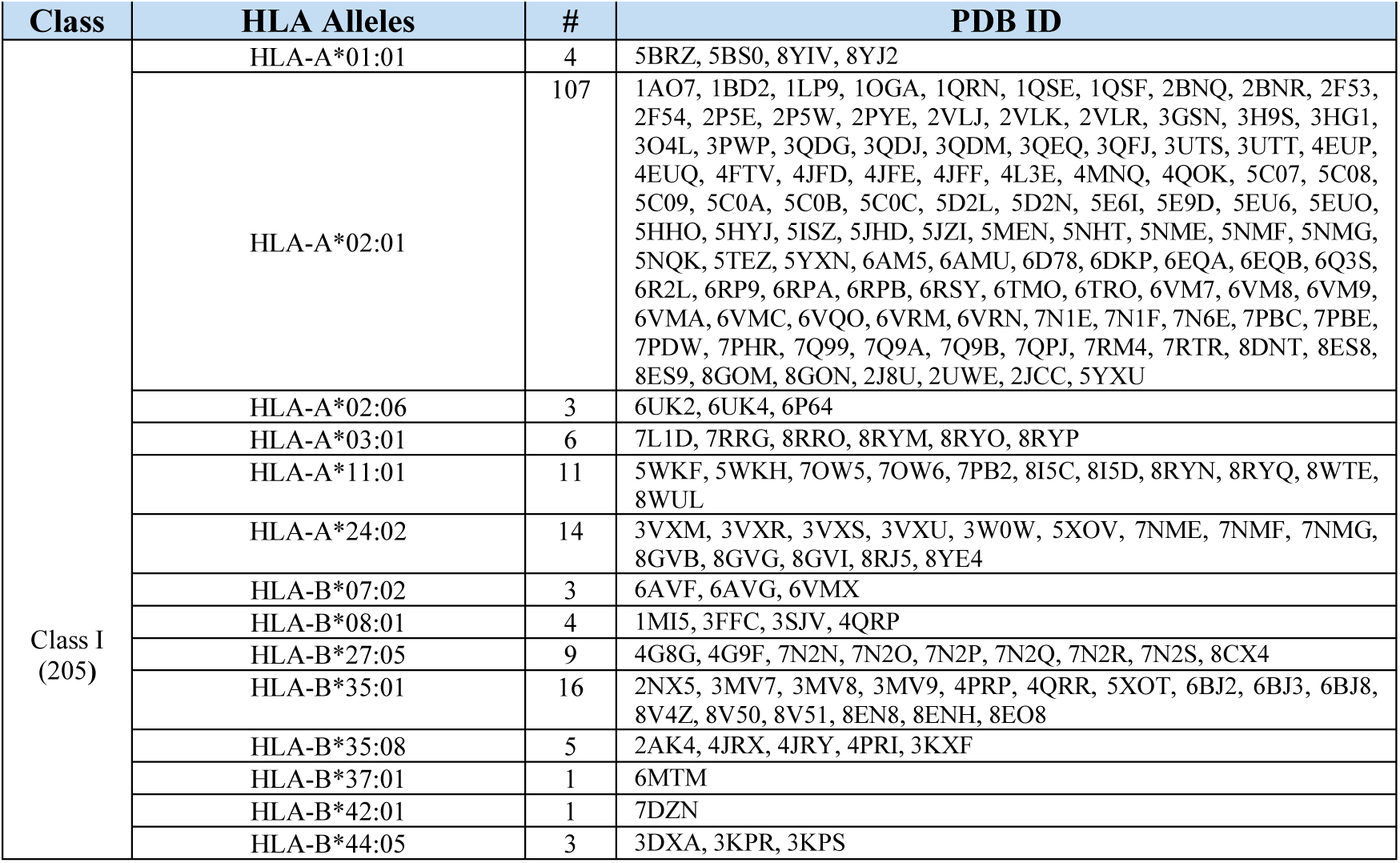
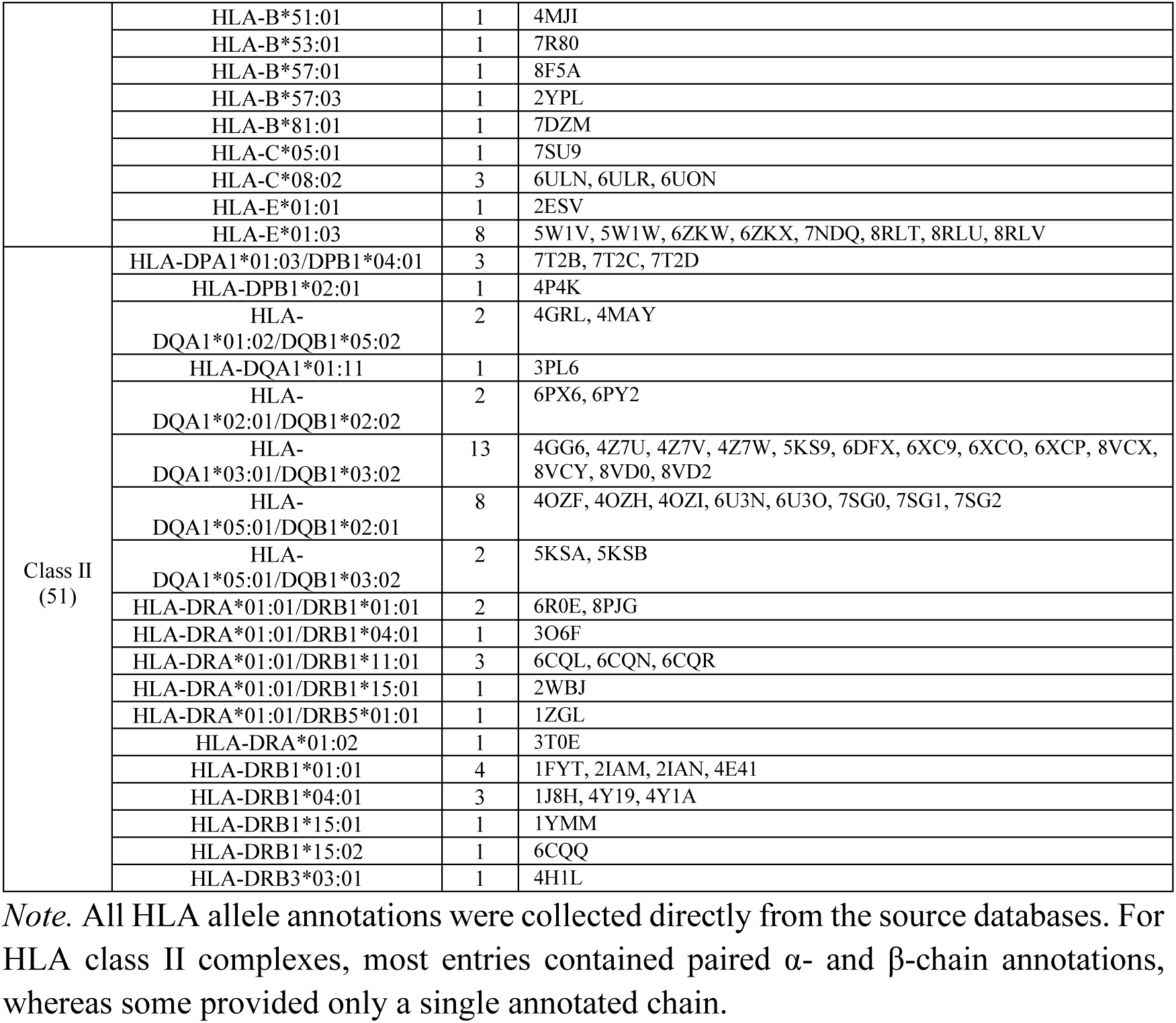
HLA classes and alleles corresponding to each PDB entry in the DynaTPH database. The “#” column indicates the total count for each HLA allele.

### Structure preprocessing

The retained complexes were subsequently subjected to necessary structural preprocessing to generate standardized TCR-pHLA monomers for MD simulations as the initial structures (Fig. 1 and Fig. S1). These preprocessing steps comprised four categories: First, for oligomeric entries, we retained a single TCR-pHLA monomer for simulation. Second, for entries in which crystal lattice asymmetry affected the biological assembly, we generated symmetric assemblies in PyMOL^36^. Third, several TCRs contained extended C-terminal helices. Since our simulations focused on the ternary interface, we removed these extended C-terminal extensions, which are outside the core TCR fold. Finally, in some PDB files, the peptide and the HLA molecule shared the same chain identifier. While this did not affect the structure coordinates, it complicated downstream analyses, so we reassigned separate chain identifiers for the peptide and HLA.

Following structural preprocessing, we obtained a final set of 256 high-quality crystal structures of TCR-pHLA ternary complexes. These curated crystal structures constitute the static structural component of DynaTPH. The dataset covers complexes involving both class I and class II HLA molecules. Table 1 summarizes the HLA classes and alleles, together with the corresponding PDB ID. They were subsequently used to initialize MD simulations, from which we generated the trajectory-based dynamic dataset and derived multidimensional physicochemical descriptors, included in the DynaTPH dataset.

### Molecular dynamic simulations

All MD simulations were performed using GROMACS version 2023.5^37^ with the CHARMM36m^38^ force field. Prior to simulation, missing residues at the TCR-pHLA binding interfaces were modeled using MODELLER^39^ to restore structural continuity and ensure physically meaningful interaction analysis. Each TCR-pHLA complex was solvated in a cubic simulation box with the complex positioned at the center. The box was aligned such that the long axis of the ternary complex was parallel to the z-axis, and a minimum distance of 15 Å was maintained between the complex and the box boundaries. Solvation was conducted using the TIP3P^40^ water model, and Na^+^ and Cl^-^ ions were added to neutralize the system at a final NaCl concentration of 0.15 M.

Energy minimization was carried out using the steepest descent algorithm until the maximum force on any atom fell below 1,000 kJ/mol/nm, ensuring the elimination of steric clashes and structural irregularities prior to equilibration. The equilibration phase consisted of two steps. First, a 1 ns simulation was performed under the canonical ensemble (NVT). Temperature was maintained at 310 K using the V-rescale thermostat^41^. Protein backbone atoms were restrained with harmonic potentials using a force constant of 1,000 kJ/mol/nm². In the second step, an additional 1 ns equilibration was performed under NPT ensemble to maintain a pressure of 1 atm. The same backbone restraints were applied, and long-range electrostatics were treated using the Particle Mesh Ewald (PME)^42^ method throughout both equilibration phases. Production simulations were conducted in the NPT ensemble for 50 ns without backbone restraints. The pressure was maintained at 1 atm using the isotropic Parrinello-Rahman barostat^43^. The integration time step remained at 2 fs, and atomic coordinates were saved every 50 ps. To enhance the repeatability of the sampling, each system was simulated three times in parallel. The TCR-pHLA structures were inspected and visualized by ChimeraX^44^ and PyMOL^36^.

### Multidimensional physicochemical data generation

To build the multidimensional physicochemical property compoent of DynaTPH and systematically characterize the TCR-pHLA recognition, multi-descriptors were generated at three hierarchical structural levels: peptide, ternary complex, and interaction interface.

Peptide Scale. We quantified residue-resolved flexibility and conformational accessibility along the presented antigen. The root mean square fluctuation (RMSF) was computed for each peptide residue to assess local conformational mobility within the HLA binding groove. The solvent accessible surface area (SASA) was calculated for all peptide atoms within the ternary complex to quantify peptide burial and solvent exposure during the simulations.

Ternary complex Scale. Global structural stability and time-dependent conformational variation of the TCR-pHLA complex were evaluated using root mean square deviation (RMSD) trajectories. RMSD values were calculated across the simulation trajectories to measure structural deviation relative to the initial conformation under a consistent simulation protocol.

Interaction-Interface Scale. Intermolecular interactions were systematically quantified at both the peptide-HLA and TCR-pHLA interfaces to characterize interface formation, rearrangement, and persistence over time. Hydrogen bonds were identified using a donor-acceptor distance cutoff of 3.5 Å to quantify interactions at the peptide-HLA interface and the TCR-pHLA interface, providing a measure of interaction stability at both binding sites. Intermolecular contact numbers were extracted using three distinct distance thresholds (3.5, 4.0 and 4.5 Å), with the final values reported as an average across these thresholds to capture interaction robustness across multiple distance scales.

Together, the data including peptide flexibility and exposure (RMSF and SASA), global structural stability (RMSD), and interface-level hydrogen bond and contact patterns, provided complementary physicochemical annotations for the MD trajectories and formed a core component of the DynaTPH database. All calculations were performed using GROMACS analysis tools^37^. Physicochemical data were generated using a consistent workflow and reported in standardized formats across systems, facilitating cross-complex comparisons and enabling downstream reuse without repeating trajectory post-processing. For each complex, dynamic descriptors were obtained from three independent replicate simulations by analyzing the final ten nanoseconds of each production trajectory and averaging values across replicates to improve consistency and reproducibility.

### Data Records

The complete DynaTPH dataset now is only available for peer review through a private Zenodo reviewer-access link. The dataset comprises curated static TCR-pHLA structures, molecular dynamics trajectories and corresponding structural frames, derived structural and biophysical features, and associated descriptor tables. A lightweight version of the dataset, excluding the dynamic data, is also available through the project GitHub repository (https://github.com/leifugroup/DynaTPH), including static structural data, extracted feature data, and associated descriptor files.

### Organization

The DynaTPH dataset is structured into three corresponding top-level directories: *Static Data*, *Dynamic Data* and *Feature Data* together with a system-level descriptors file *descriptor.csv* and a residue-level dynamic descriptors file *rmsf.csv*. A schematic overview of the database organization and file paths is shown in Fig. 3.

**Fig. 3.**
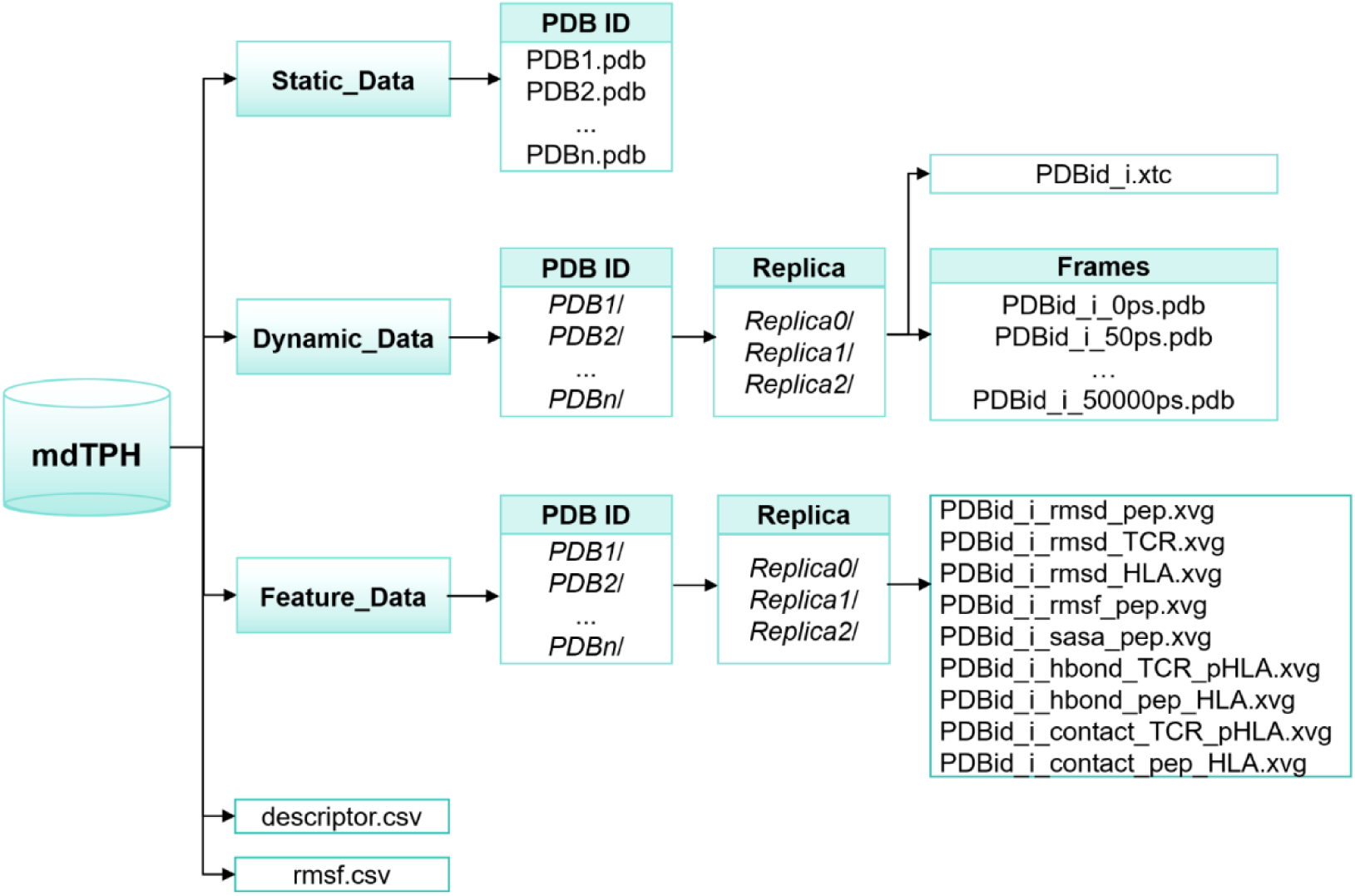
Schematic diagram of the DynaTPH dataset structure.

The *Static Data* directory contains curated TCR-pHLA complex structures used as initial configurations for molecular dynamics simulations. Each structure is provided in PDB format and named by its PDB ID, resulting in a total of 256 complexes.

The *Dynamic Data* directory stores molecular dynamics trajectories and corresponding structural frames. It contains 256 subfolders organized by PDB ID (e.g., 1BD2). Each complex folder includes three replicate simulations (*Replica0*, *Replica1*, and *Replica2*), corresponding to three independent MD simulations. Within each *Replica* folder, the integrated 50 ns trajectory files in “xtc” format and the corresponding *Frames* directory are provided. The *Frames* directory contains structures extracted from 50 ns trajectories at 50 ps intervals, resulting in 1001 frames per replica. The first frame (0 ps) corresponds to the initial TCR-pHLA complex structure. Throughout the dataset, The index *i* in file names corresponds to *Replica0*, *Replica1*, and *Replica2* (*i* = 0, 1, and 2), respectively. All trajectories and frames include only the TCR-pHLA complex, with solvent molecules and ions removed.

The *Feature Data* directory contains multidimensional physicochemical property data analyzed from simulated trajectories. Each PDB ID directory contains replica-specific subfolders structured similarly to those in the *Dynamic Data* directory. Each *Replica* folder includes multidimensional physicochemical data such as RMSD (peptide, TCR, and HLA), peptide RMSF and SASA, as well as hydrogen bond and contact profiles for both peptide-HLA and TCR-pHLA interfaces. In the file naming scheme, *TCR_pHLA* denotes the TCR-pHLA interface and *pep_HLA* denotes the peptide-HLA interface.

The system-level descriptors file *descriptor.csv* organized by PDB ID, provides a comprehensive summary of the structural, physicochemical, and dynamic properties of each TCR-pHLA complex, with each entry indexed by PDB ID. It includes HLA classes and alleles, peptide length, amino acid sequence, as well as selected physicochemical properties such as peptide SASA, and hydrogen bond and contact numbers at the peptide-HLA and TCR-pHLA interfaces. These values were averaged across three independent trajectories using a consistent analysis workflow. And the structure preprocessing notes applied during the structure preparation prior to molecular dynamics simulations were also recorded in this file.

The residue-level dynamic descriptors file *rmsf.csv* describes the RMSF of each peptide residue in each TCR-pHLA complex.

### Size

The DynaTPH database contains 256 curated TCR-pHLA ternary complex structures in PDB format within the *Static Data* folder. Among them, there are 205 HLA class I structures and 51 HLA class II structures (Fig. 4A). Fig. 4B and 4C summarize the distributions of residue counts and atom counts across these complexes. Most systems contain more than 800 amino acid residues and approximately 13,000 atoms when considering the TCR-pHLA ternary complex only. In the corresponding solvated molecular dynamics systems, the inclusion of explicit water molecules and ions increases the system size to approximately 500,000 atoms per simulation system. The *Dynamic Data* folder comprises molecular dynamics trajectories for all 256 complexes, each simulated in three independent replicas, resulting in a total of 768 trajectories (256 × 3). Each trajectory contains 1,001 structural frames extracted at 50 ps intervals from a 50 ns simulation, yielding 768,768 frames in total. The cumulative simulation time across all systems is 38.4 μs (256 × 3 × 50 ns). The *Feature Data* folder contains per-trajectory physicochemical descriptors, including RMSD, RMSF, SASA, hydrogen bond, and contact numbers. For each complex, nine descriptor files are provided per replica, resulting in a total of 6,912 files (256 × 3 × 9). Overall, the DynaTPH database occupies approximately 800 GB of storage (Fig. 4D).

**Fig. 4.**
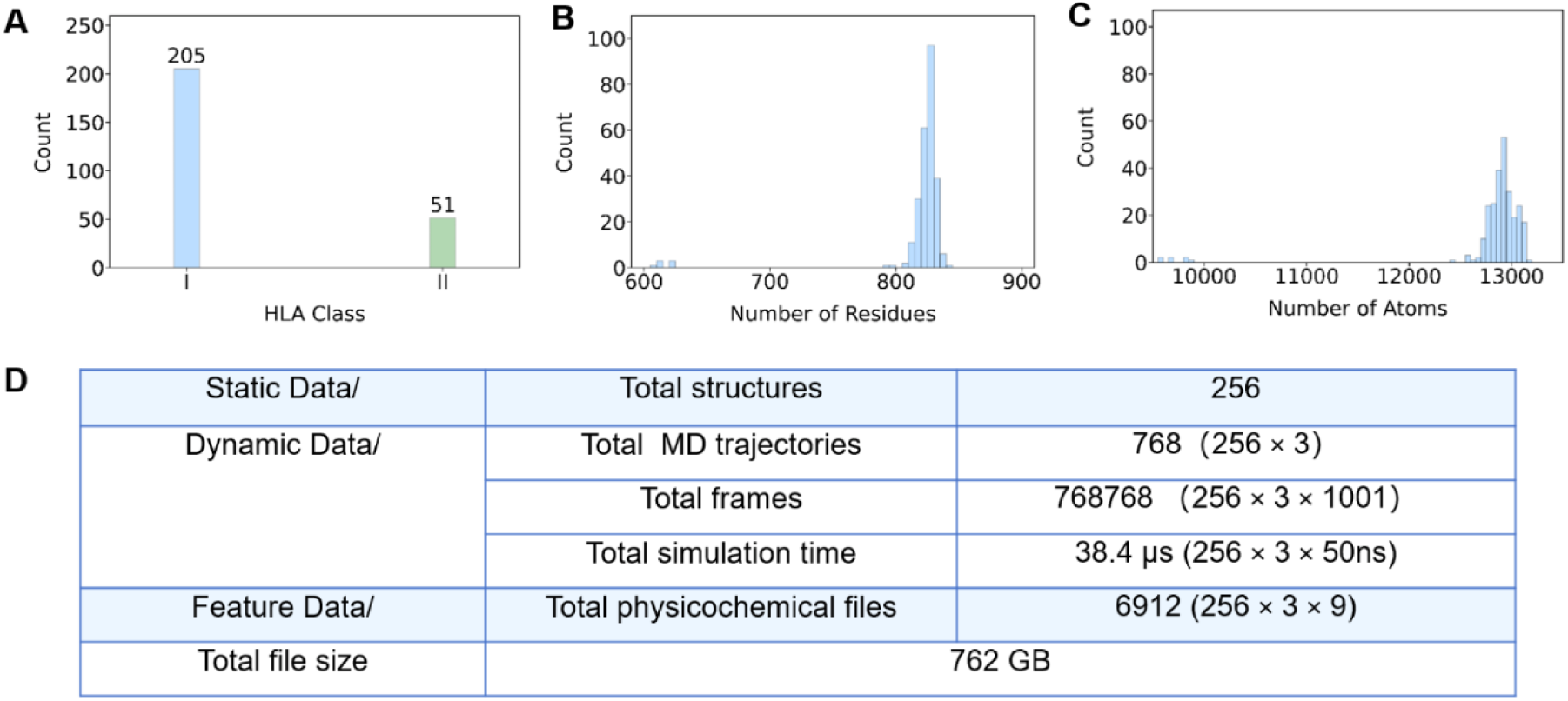
The DynaTPH database statistical overview. (A) HLA class distribution. (B) Distribution of residue numbers in TCR-pHLA complexes. (C) Distribution of atom numbers in TCR-pHLA complexes. (D) Summary statistics of the dataset.

## Summary

### Molecular diversity of the DynaTPH dataset

The DynaTPH dataset comprises a broad collection of currently resolved TCR-pHLA complex structures and captures substantial molecular diversity in HLA classes, HLA alleles, and peptide length. To characterize the molecular heterogeneity, we analyzed the distribution of HLA classes and alleles represented in the DynaTPH. Among all complexes, 205 structures (80%) are associated with HLA class I molecules, while 51 structures (20%) correspond to HLA class II molecules (Fig. 4A). The HLA class I group includes HLA-A, HLA-B, HLA-C, and HLA-E genes, whereas HLA class II structures comprise HLA-DP, HLA-DR, and HLA-DQ genes. In total, 42 distinct HLA alleles are present in the dataset (Table 1). Among these, HLA-A*02:01 is the most frequently observed allele, accounting for 107 complexes (Fig. 5A).

**Fig. 5.**
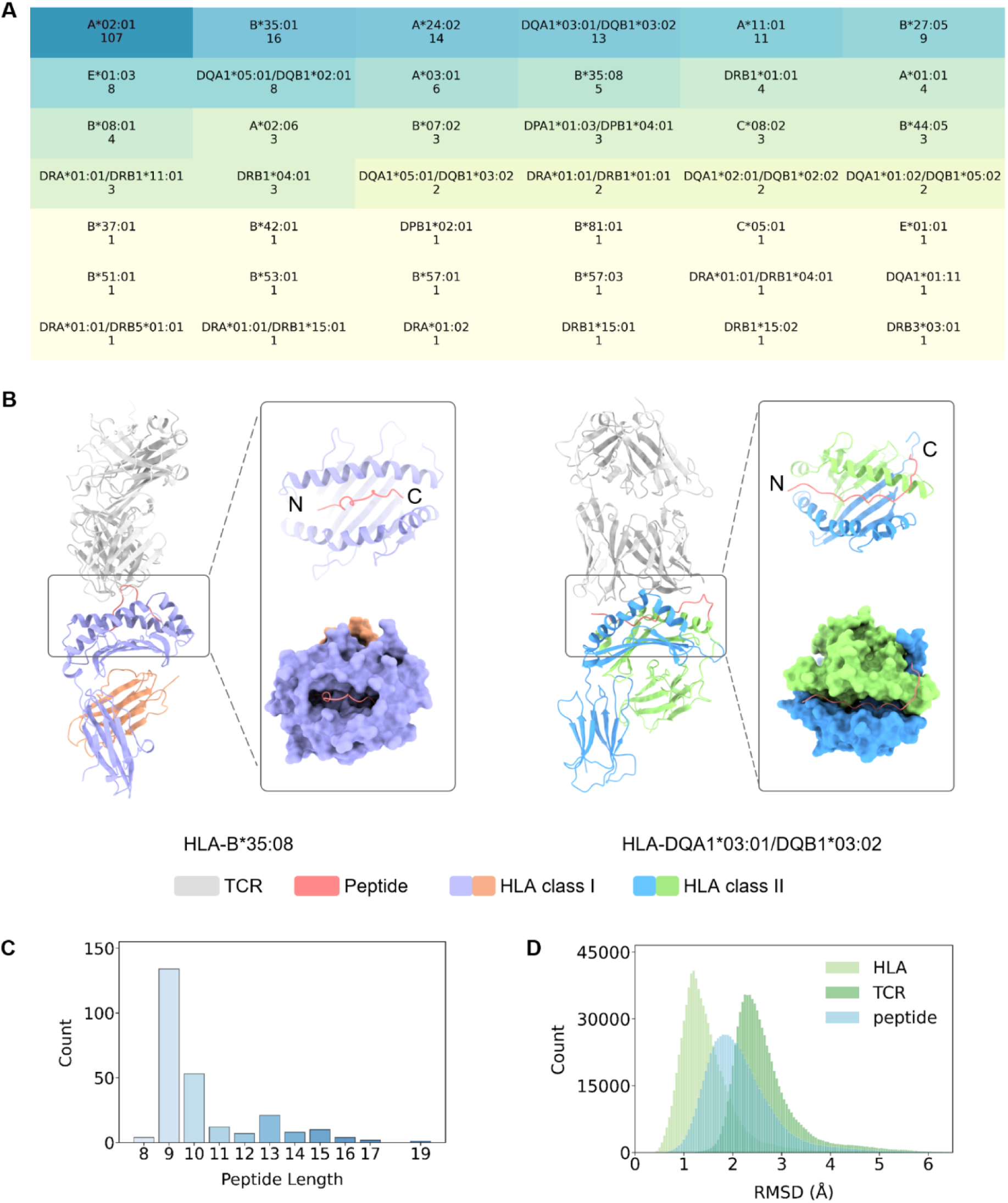
Composition and structural stability of the DynaTPH dataset. (A) Heatmap of HLA alleles in the dataset, ranked by frequency. Omit the prefix “HLA-”. (B) Class I complex (left, PDB: 2AK4, 13-mer peptide) and Class II complex (right, PDB: 6DFX, 19-mer peptide). Zoomed-in views show peptide-HLA interfaces. (C) Peptide length distribution. (D) Distribution of the RMSD of protein heavy atoms across all TCR-pHLA complexes between the first and last frames of the simulation trajectory for and all replicas. HLA (light green), TCR (green), and peptide stability relative to the HLA molecule, peptide (blue).

The differences in peptide length distributions reflect the distinct structural characteristics of HLA class I and class II binding grooves. HLA class I molecules possess short, closed binding grooves that typically accommodate shorter peptides, peptides longer than approximately ten residues often adopt a solvent-exposed central bulge. In contrast, HLA class II molecules have binding grooves that are open at both ends, allowing longer peptides to bind and extend beyond the groove (Fig. 5B). To quantify these structural differences, we analyzed peptide length distribution of across all complexes. Peptide lengths range from 8 to 19 amino acids, with short 9-mer peptides being the most abundant (100 entries, Fig. 5C), consistent with the predominance of HLA class I complexes in the dataset. In the DynaTPH dataset, peptides presented by HLA class I molecules span 8-13 residues, whereas those bound to HLA class II molecules span 13-19 residues (see descriptors.csv). These peptide length distributions are consistent with previously reported binding preferences of HLA class I and class II molecules.

We further evaluated global structural properties of the TCR-pHLA complexes, including total residue counts and atom numbers (Fig. 4B, C). The distributions indicate that most complexes contain more than 800 amino acid residues and approximately 13,000 atoms in the TCR-pHLA ternary structure, without evident extreme outliers. These statistics suggest that the structures are consistently processed while maintaining biologically relevant molecular variability. These results demonstrate that DynaTPH captures diverse TCR-pHLA compositions across HLA classes, alleles, and peptide lengths in a systematically curated manner.

### Structural stability of TCR-pHLA complexes

To evaluate the reliability of the MD trajectories in DynaTPH, we computed and summarized RMSD distributions across all independent replicate simulations (Fig. 5D). Three RMSD metrics were analyzed: (i), HLA, (ii) TCR, and (iii) peptide. For the peptide, RMSD was calculated relative to the HLA molecule to characterize the stability and dynamic behavior of the peptide within the HLA binding groove. Across all systems and replicas, RMSD distributions were highly consistent. HLA molecules exhibited the lowest RMSD values, consistent with high structural stability of the HLA scaffold. The peptide RMSD relative to HLA was modestly higher, indicating that peptide binding was maintained while allowing local conformational fluctuations within the groove. TCR molecules displayed higher RMSD, reflecting their comparatively greater conformational flexibility during pHLA recognition. Importantly, although the overall TCR-pHLA binding architecture was preserved throughout the simulations, all systems retained measurable conformational variability across the trajectories. This indicates that the simulations capture dynamically fluctuating ensembles rather than static structures. The reproducibility of RMSD distributions across independent replicas further supports that the trajectories reached comparable equilibrated states while preserving biologically relevant molecular motions (Fig. S2). RMSD analyses show that DynaTPH provides structurally stable yet dynamically rich conformational ensembles suitable for downstream studies of TCR-pHLA recognition, interfacial interactions, and peptide dynamics.

### Interactions in TCR-pHLA complexes

We analyzed hydrogen bonds and contacts at the peptide-HLA and TCR-pHLA interfaces to assess whether DynaTPH captures physically plausible and biologically consistent interaction patterns across systems (Fig. 6A, B). Both interfaces displayed consistent interaction distributions across the dataset. The peptide-HLA interface exhibited generally higher numbers of hydrogen bonds and contacts than the TCR-pHLA interface, reflecting the stable anchoring of peptides within the HLA binding groove (Fig. S3). In contrast, interactions between TCRs and pHLA complexes involved fewer contacts and hydrogen bonds, consistent with the comparatively greater flexibility of TCR engagement. These trends were observed for both HLA class I and class II complexes, in agreement with prior structural and biophysical studies^45–48^. Notably, hydrogen bond numbers varied within a relatively narrow range across systems, whereas contact numbers spanned a broader range. This difference reflects the distinct interaction features represented by these metrics, with hydrogen bonds representing specific and directional interactions, whereas contacts reflect broader interfacial engagement. Collectively, these interaction distributions demonstrate that DynaTPH captures structurally plausible and biologically consistent TCR-pHLA interaction patterns, supporting the reliability of the dataset for downstream analyses.

**Fig. 6.**
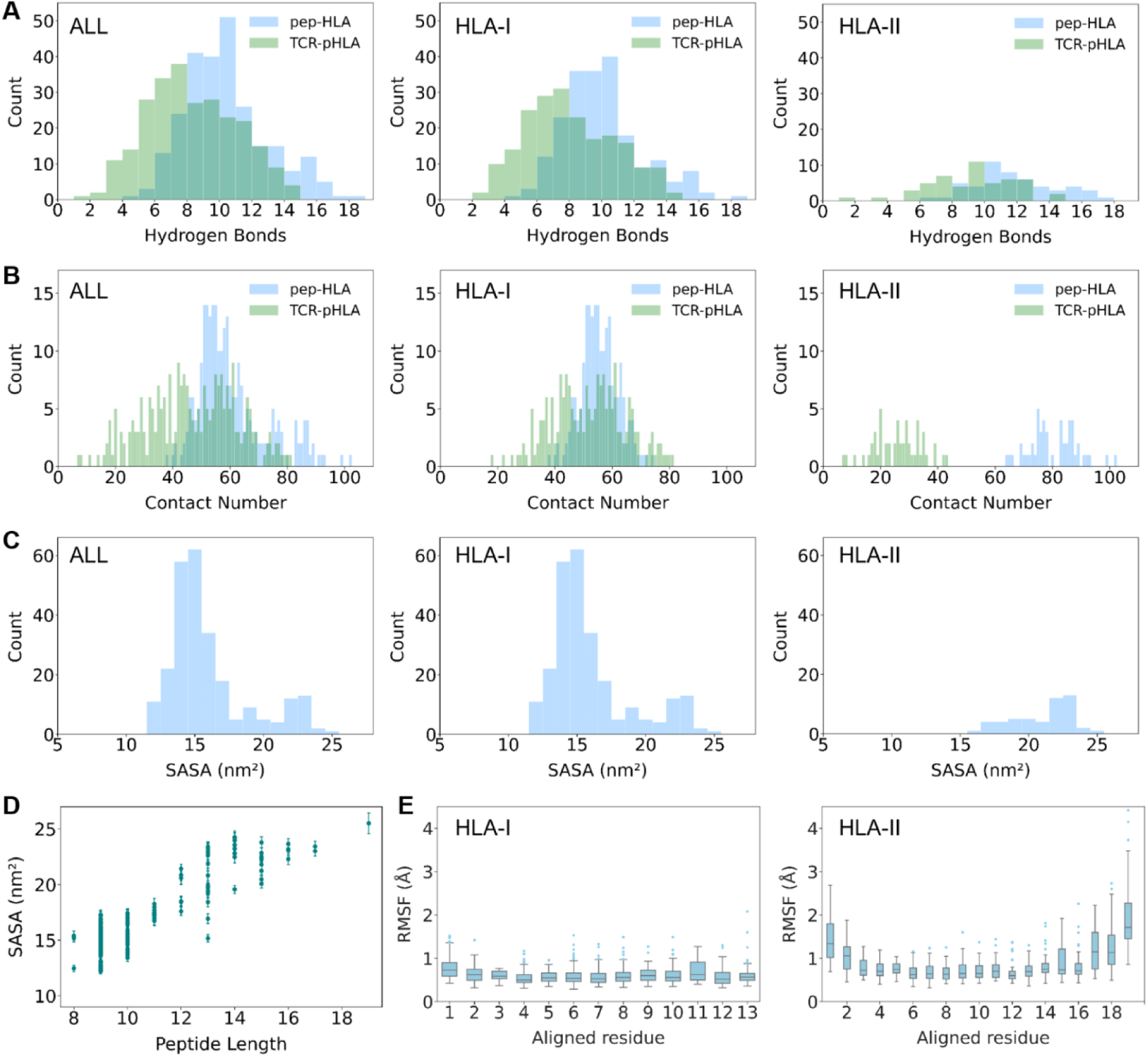
Interaction characteristics and peptide dynamics in TCR-pHLA complexes. (A) Hydrogen bond distributions at the peptide-HLA (blue) and TCR-pHLA (green) interfaces across the full dataset, HLA-I, and HLA-II. (B) Contact distributions for peptide-HLA and TCR-pHLA. (C) SASA distributions of peptide. (D) Relationship between peptide SASA and peptide length. (E) RMSF of peptide aligned residues, presented separately for HLA-I and HLA-II.

### Peptide flexibility and solvent exposure

We characterized peptide dynamics at the binding interface using RMSF and SASA (Fig. 6C, D). Peptide SASA ranged from 12 to 26 nm^2^, with most values falling within a limited range (Fig. 6C), reflecting the constrained solvent exposure imposed by peptide binding within the HLA groove. Peptides bound to HLA-II had larger SASA than those bound to HLA-I, consistent with the open-ended binding groove of class II molecules that accommodates longer peptides extending beyond the groove (Fig. 6C). Consistent with this trend, SASA showed a positive correlation with peptide length, as longer peptides typically exhibited greater solvent exposure (Fig. 6D).

Peptide flexibility was quantified by calculating the RMSF for each peptide residue and averaging it across three replicas. To enable comparisons among peptides of different lengths, peptides within each HLA class were aligned by sequence position and padded to the maximal length observed in that class (13 residues for HLA-I and 19 residues for HLA-II). RMSF values were then averaged at each aligned position (Fig. 6E). RMSF results showed that longer peptides exhibited higher flexibility, which was positively associated with solvent exposure, indicating that more exposed regions tend to be more dynamic. Clear differences in flexibility were observed between HLA classes. Peptides bound to HLA-I displayed lower overall flexibility, whereas HLA-II associated peptides showed increased flexibility at terminal regions, with comparatively restricted fluctuations in the central region.

These flexibility and exposure patterns are consistent with known structural features of HLA peptide binding, including the closed-ended groove of HLA-I molecules and the open-ended groove of HLA-II molecules. Together, these SASA and RMSF trends support the ability of DynaTPH to capture realistic dynamics and interfacial behavior across diverse TCR-pHLA complexes.

## Code Availability

Custom scripts used for data collection and screening of the DynaTPH dataset are available through the project GitHub repository (https://github.com/leifugroup/DynaTPH). Molecular dynamics simulations and subsequent trajectory analyses were performed using the GROMACS 2023 package^37^ and standard GROMACS analysis tools.

## Funding

This study was funded by the National Key Research and Development Program of China (2025YFF0515300); National Natural Science Foundation of China (62425112); Shenzhen Medical Research Fund (E250200620, E250200622, E250200623); Shenzhen Key Laboratory Construction Project (SYSRD20250529113406008).

## Notes

### Competing Interest Statement

The authors have declared no competing interest.

