## supplementary for "A multi-scale structural and biophysical atlas of TCR-peptide-HLA recognition dynamics"

**Table S1. Comparison of publicly available TCR-related databases and resources reviewed in this study.**

| Database | TCR sequence | Antigen specificity | HLA annotation | Structural information* |
|---|---|---|---|---|
| STCRDab[1] | √ | √ | √ | √ |
| TCR3d[2] | √ | √ | √ | √ |
| TRAIT[3] | √ | √ | √ | √ |
| IEDB[4] | Partial | √ | √ | Partial |
| VDJdb[5] | √ | √ | √ | Partial |
| ATLAS[6] | √ | √ | √ | Partial |
| TBAdb[7] | √ | √ | √ | × |
| McPAS[8] | √ | √ | Partial | × |
| TCRdb[9] | √ | × | × | × |
| NeoTCR[10] | √ | √ | √ | × |

*Structural information denotes experimentally determined TCR-pHLA structures or links to corresponding structural records. Partial indicates availability for only a subset of entries.

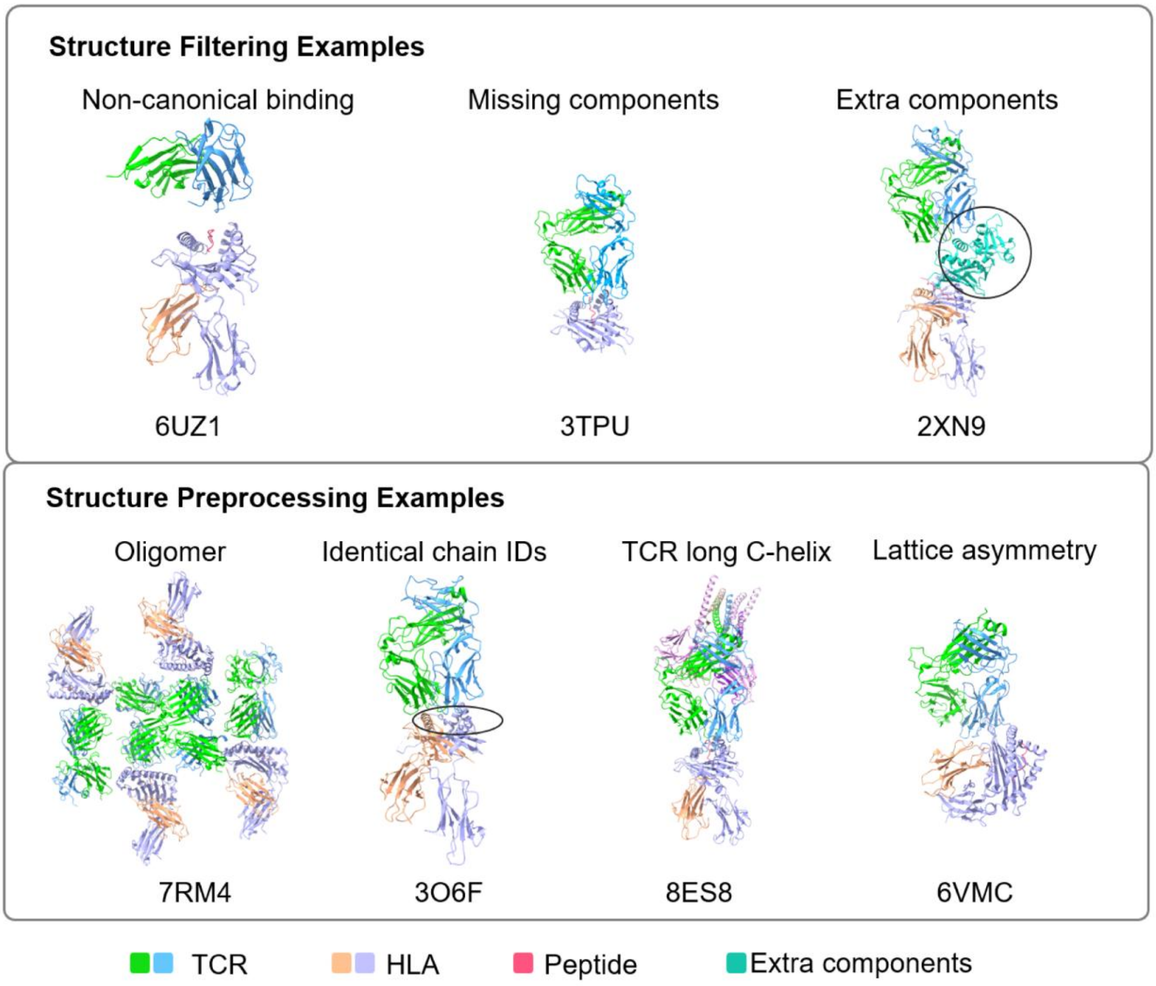


**Fig. S1. Representative examples of structural filtering and preprocessing of TCR-pHLA complexes.** Examples of structures excluded during manual quality control are shown in the upper panel, including non-canonical TCR-pHLA binding orientations (PDB: 6UZ1), missing structural components (PDB: 3TPU), and complexes containing extra components beyond the canonical TCR-pHLA assembly (PDB: 2XN9; circled region). The lower panel shows representative preprocessing operations, including extraction of biologically relevant monomers from oligomeric complexes (PDB: 7RM4), reassignment of duplicated chain identifiers (PDB: 3O6F; circled region), where the peptide and one HLA chain initially shared the same chain identifier and were therefore displayed in the same color prior to correction, removal of long TCR C-terminal extensions (PDB: 8ES8), and reconstruction of biologically relevant complexes from crystallographic asymmetric units (PDB: 6VMC).

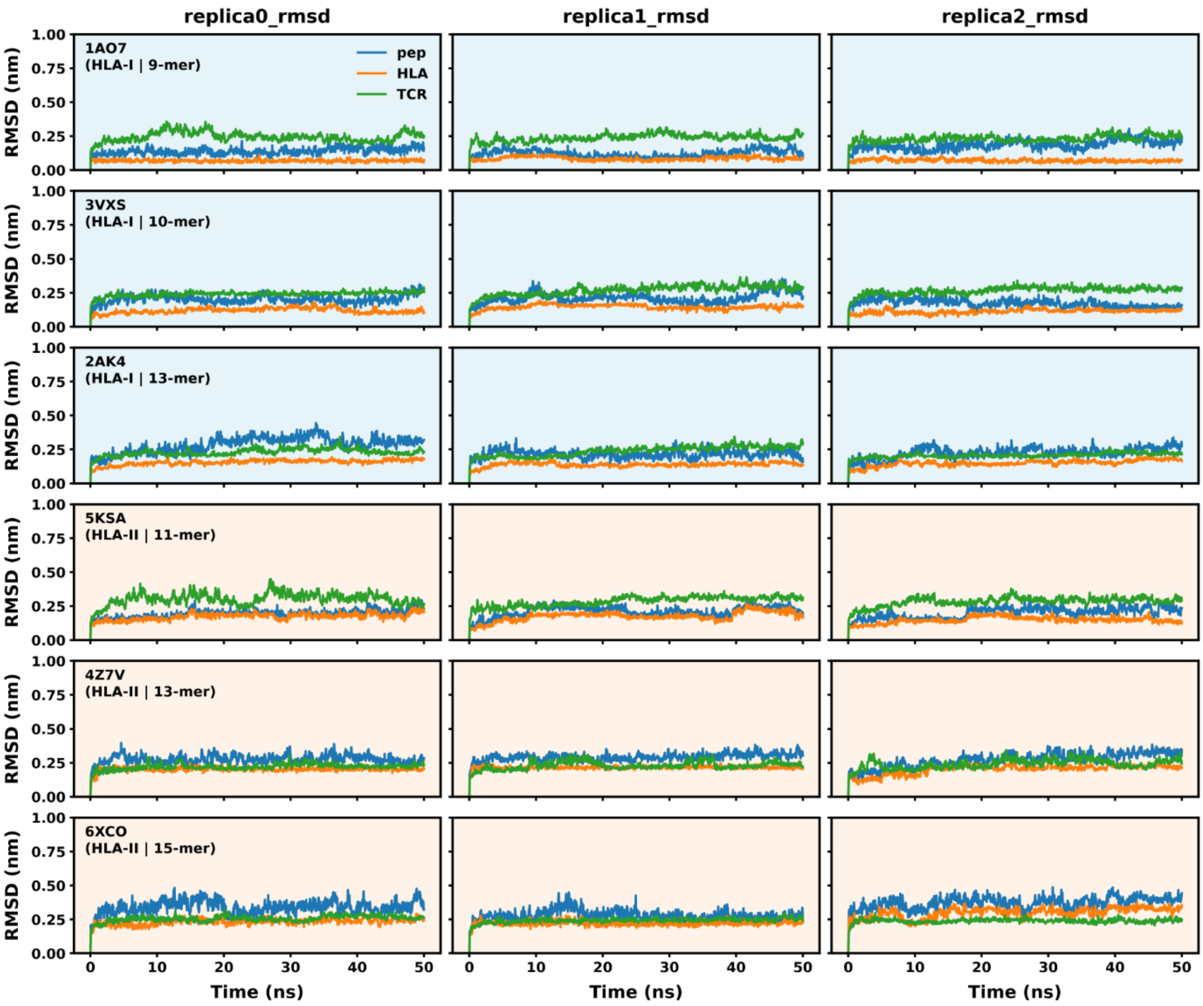


**Fig. S2. Representative RMSD profiles of TCR-pHLA complexes with different peptide lengths and HLA classes.** RMSD profiles of three independent molecular dynamics replicas for representative TCR-pHLA complexes containing peptides of different lengths. The upper three rows (light blue background) represent HLA class I complexes, and the lower three rows (light orange background) represent HLA class II complexes. Peptide lengths are indicated in the corresponding panels. The RMSD trajectories of the peptide, HLA, and TCR are shown in blue, orange, and green, respectively. All systems reached a stable conformational state during the simulations.

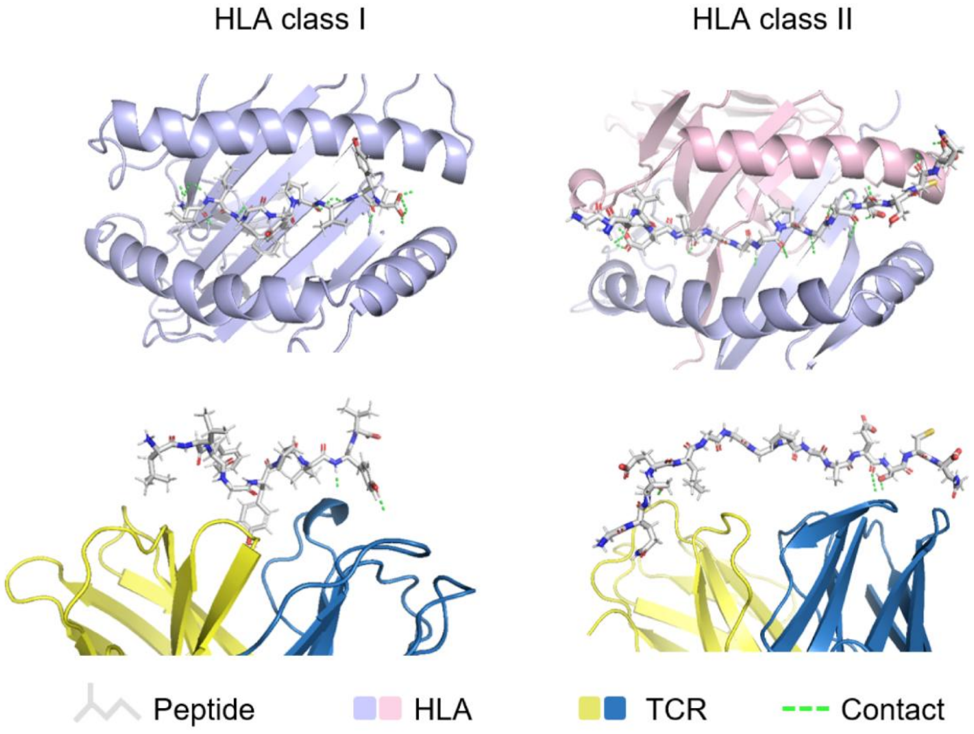


**Fig. S3. Representative peptide-HLA and peptide-TCR contacts in HLA class I and class II TCR-pHLA complexes.** Contact snapshots for an HLA class I complex (PDB: 1BD2; left) and an HLA class II complex (PDB: 8VCX; right). The first row shows top-view snapshots of peptide-HLA contacts, whereas the second row shows peptide-TCR contacts. Peptides are shown as sticks, while TCR and HLA molecules are shown as cartoons. Green dashed lines indicate contacts identified using a 5 Å distance cutoff. In both representative complexes, the peptide forms more extensive contacts with HLA than with TCR.
